# Durable Control of HIV-1 Using a *Staphylococcus aureus* Cas9-Expressing Lentivirus Co-Targeting Viral Latency and Host Susceptibility

**DOI:** 10.1101/2020.08.10.243329

**Authors:** Leonard R. Chavez, Nishith R. Reddy, Kyle A. Raymond, Mohamed S. Bouzidi, Shivani Desai, Zain Y. Dossani, Hannah S. Sperber, James Theiler, Bette Korber, Satish K. Pillai

## Abstract

CRISPR/Cas9 gene editing has the potential to revolutionize the clinical management of HIV-1 infection, and may eliminate the need for antiretroviral therapy (ART). Current gene therapies attempt to either excise HIV-1 provirus or target HIV-1 entry receptors to prevent infection of new cells. Using a viral dynamic model, we determined that combining these two interventions, in the presence or absence of ART, significantly lowers the gene editing efficacy thresholds required to achieve an HIV-1 cure. To implement this dual-targeting approach, we engineered a single lentiviral vector that simultaneously targets multiple highly-conserved regions of the provirus and the host CXCR4 coreceptor, and developed a novel coculture system enabling real-time monitoring of latent infection, viral reactivation, and infection of new target cells. Simultaneous dual-targeting depleted HIV-1-infected cells with significantly greater potency than vectors targeting either virus or host independently, highlighting its potential as an HIV-1 cure strategy.

## INTRODUCTION

Antiretroviral therapy (ART) has been successful in suppressing HIV-1 replication in infected individuals, reducing virus-associated mortality and morbidity^1^. However, viral eradication cannot be achieved with ART due to the persistence of a reservoir of latently infected cells that harbor replication-competent virus^2,3^. Therefore, multiple therapeutic approaches are now focused on either preventing or delaying viral rebound following treatment interruption, resulting in a functional cure that would be characterized by a long-term remission period. Current strategies include immunotherapy and vaccination to prevent latent reactivation or to reduce viremia following reactivation^4,5^, reactivation of latent provirus, also known as the shock and kill approach^6^, and gene therapy to either target the provirus directly or make cells refractory to HIV-1 infection^7-19^.

CRISPR/Cas9 gene-editing technology has given rise to the hope of achieving a functional cure through gene therapy. Multiple pre-clinical studies have already demonstrated the potential of using CRISPR/Cas9 to directly excise/disrupt the HIV-1 provirus^7-15^, and to prevent the infection of new cells via knockout of HIV-1 co-receptors^16-19^. However, achieving a functional cure through either of these strategies alone is not currently feasible.

HIV-1 cure through direct targeting of the provirus presents several challenges. The first problem is the vast diversity of circulating HIV-1 strains within and among HIV-1-infected individuals, and the need to account for this diversity when identifying potential guide RNA (gRNA) target sites within the HIV-1 provirus. The second problem arises from the observation that CRISPR/Cas9 gene editing of the provirus can lead to escape mutants via non-homologous end joining repair (NHEJ)^20-24^. These two problems can, partially, be solved by targeting multiple conserved sites, since the occurrence of multiple mutations is less likely than the occurrence of one, and targeting more conservative regions of the provirus leads to a delay in the generation of escape variants^24^. Unfortunately, this solution gives rise to a third problem: the requirement of efficiently delivering multiple gRNAs and a Cas9 enzyme into target cells *in vivo*. Lentiviral vectors are best suited to overcome this barrier due to their large cargo capacity and low antigenicity. However, they suffer—as do all current *in vivo* gene delivery methods— from the inability to achieve adequate transduction efficiency. This problem is amplified by the inference that 99.9% of infected cells would need to be successfully edited in order to effectively curb the latent reservoir^25^.

Alternatively, rendering cells refractory to HIV-1 infection via *CCR5* or *CXCR4* editing benefits from an *ex vivo* approach, but is still not devoid of obstacles. First and foremost, is the requirement that >88% of cells need to be made refractory to infection in order to prevent viral propagation, though lower efficiency may still be beneficial^25^. Additionally, this approach completely ignores the population genetics of latent virus, leaving open the possibility for the virus to reactivate and use alternative co-receptors to enter target cells and initiate expansion.

Despite these challenges, achieving long-term remission, and thus a functional cure, is attainable, and the blueprint to accomplish this can be found in the case studies of both the Berlin and London patients^26-28^. In receiving bone marrow transplants from CCR5-deficient donors, these patients were treated with a combined approach: First, transplant conditioning shrunk the latent reservoir by removing infected cells; and, second, the transplant itself replaced cells susceptible to infection with resistant cells. Thus, targeting both provirus and co-receptor concomitantly constitutes a successful two-pronged approach that may lower the threshold of edited cells required to produce a long-term remission period within infected individuals.

Here, we report a proof of concept study demonstrating the use of CRISPR/Cas9 gene-editing technology to simultaneously target HIV-1 provirus in latently infected cells, and render uninfected cells non-permissive to infection through co-receptor editing. We use a dynamic mathematical model (see Methods) to simulate antiviral genome editing strategies targeting viral latency and host susceptibility, predicting that gene editing efficacies required to achieve long-term remission are significantly reduced when these interventions are combined. Importantly, the advantage of co-targeting is evident both when gene therapies are administered during suppressive ART, and in the absence of ART when administered in the setting of viremia. We then simultaneously target viral latency and host susceptibility using a single all-in-one lentiviral vector that expresses *Staph. aureus* Cas9 (SaCas9), two gRNAs targeting highly conserved regions of the HIV-1 genome, and a single gRNA targeting *CXCR4*. By designing and implementing a novel *in vitro* co-culture assay, we demonstrate successful multiplex editing using our single lentiviral vector. Further, we show that simultaneous CRISPR/Cas9 targeting of provirus and co-receptor results in significantly lower frequencies of infected cells as compared to strategies targeting either alone.

## RESULTS

### Mathematical modeling predicts co-targeting of latent provirus and co-receptor reduces the threshold efficacy required to achieve long-term remission

The use of a bispecific CRISPR/Cas9 vector capable of simultaneously targeting latent provirus and co-receptor could both reduce virion production from infected cells, and the fraction of target cells susceptible to infection (Figure 1A-B). To estimate the minimum efficacy required in this two-pronged approach to achieve a functional cure, we used a mathematical model of within-host HIV-1 infection and adapted parameters and variables from previous studies^25,29-31^. Using our model, we considered the case where gene therapy is administered during concomitant ART (Figure 1A, Scenario 1). In accordance with previous reports^25^, our model estimates that these interventions in a mono-therapeutic setting would result in a functional cure only if provirus is targeted in >99% of infected cells, associated with ∼88% reduction in virion production (Supplementary Figure 1E), or if the pool of susceptible cells was diminished by >88% (Figure 1C).

**Figure 1.**
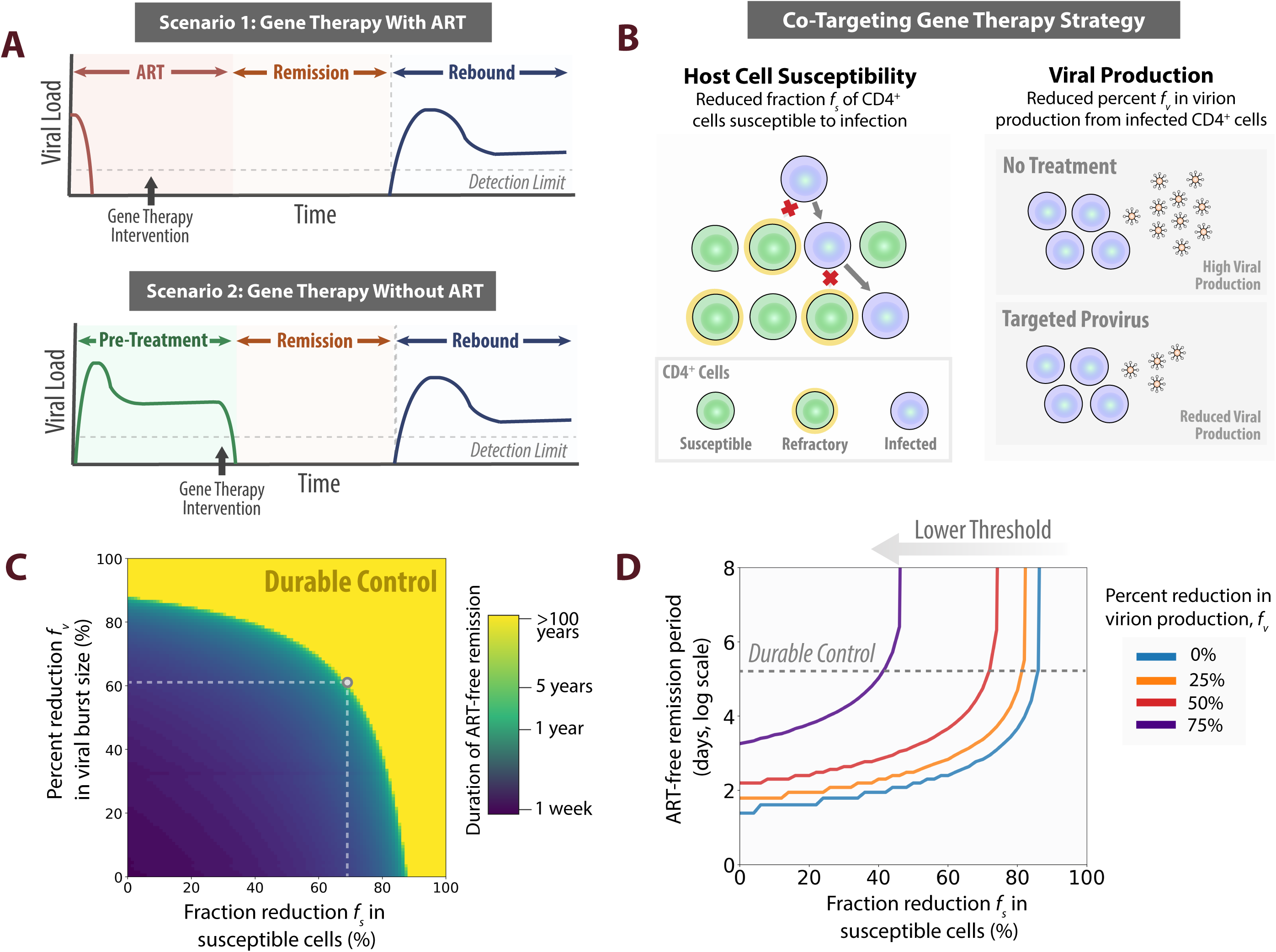
Computational predictions of within-host HIV-1 dynamics for gene therapy co-targeting host susceptibility and viral production. (A) HIV-1 gene therapy administration frameworks. Scenario1: If gene therapy interventions are administered during continuous antiretroviral therapy (ART), we define the remission period as the duration of time post-cessation of ART with viral load below clinically detectable levels *V*(*t*) ≤ 50. Scenario 2: If gene therapy is administered in the absence of ART, we define the remission period as the duration after gene therapy with viral load below clinically detectable levels. (B) Co-targeting gene therapy strategy. Using a modified standard model of in-host HIV-1 dynamics (see Methods), we model gene therapy targeting host susceptibility as a reduced fraction *f*_*s*_ of CD4^+^ cells susceptible to infection and gene therapy targeting viral production as a percent reduction *f*_*v*_ in virion production from infected CD4^+^ cells. (C) ART-free remission period with co-targeting gene therapy. Co-targeting gene therapies reduce the threshold efficacy of gene therapy needed to achieve durable control (>100 years remission) relative to single-targeting therapies. The ART-free remission period is shown for Scenario 1. (D) Lower threshold for functional cure with co-targeting gene therapy strategy. Co-targeting therapies shift the threshold needed to achieve a functional cure. The ART-free remission period is shown for Scenario 1.

Our modeling suggests that a dual-targeting gene therapy strategy can achieve a functional cure at much lower efficacies. For example, interventions which both reduce virion production and the susceptible cell population by 60% and 68%, respectively, are expected to achieve indefinite, long-term remission (Figure 1C). Targeting both infected cells and susceptible cells can lower the threshold efficiency required for durable control of infection and shift the threshold into a therapeutically accessible range (Figure 1D).

Using our viral dynamic model, we also consider what efficacy of gene therapy is required to achieve durable control in the complete absence of ART (Figure 1, Scenario 2; Supplementary Figure 1A). Our model suggests that durable control can be achieved by a bispecific gene therapy in the complete absence of ART at similar threshold efficacies as in the presence of ART (Supplementary Figure 1D). At gene therapy efficacies lower than the cure threshold, it is predicted that concomitant ART at the time of gene therapy administration can extend the expected duration of the post-treatment remission period, in proportion to ART efficacy (Supplementary Figure 1B). Furthermore, a combination of gene editing and ART efficacy affects the transition time to suppress viral load below the clinical detection limit after the administration of the gene therapy intervention (Supplementary Figure 1C). Our modeling of within-host viral dynamics in the context of this two-pronged gene therapy approach suggests that this combination strategy is a promising therapeutic intervention to achieve a functional cure both in the presence or absence of ART.

### Adapted HIV-1 vaccine immunogen design algorithm identifies highly-conserved gRNA target sites within the HIV-1 provirus

In order to test this dual gene editing approach, we first wanted to identify gRNA target sites that are as conserved as possible among HIV-1 M-group subtypes. We identified the most highly-conserved SaCas9 gRNA target sites across the HIV-1 genome using the Los Alamos National Lab (LANL) HIV Sequence Database (http://www.hiv.lanl.gov/). We interrogated the forward and reverse strands of full-length HIV-1 sequences obtained from 3,263 infected individuals, identified 400 gRNAs with the best potential for targeting HIV-1 provirus via SaCas9 based on retention of the PAM (NNGRR) sequence, and ranked the 21 base gRNA sequence preceding the PAM sequence by matches to natural HIV-1 variants. For a gRNA+PAM sequence to be considered a match, we required that the 10 bases proximal to the PAM sequence be a perfect match, and we allowed a single mismatch in the 11 most distal bases. The 20 best candidates were considered for experimental evaluation..

### High-throughput screening of candidate gRNAs targeting the provirus identified five gRNAs capable of editing HIV-1 provirus in latent cells

In order to test the candidate gRNAs identified using our bioinformatic approach for their ability to edit the HIV-1 provirus, we utilized an *in vitro* high-throughput screening assay in J-Lat cells that contain a transcriptionally silent, but reactivatable HXB2 laboratory strain of HIV^32^ (Figure 2A). Individual candidate gRNAs were *in vitro-*transcribed, and nucleofected into J-Lat cells stably expressing SaCas9. 48hrs post nucleofection, genomic DNA was isolated and the targeted proviral regions were evaluated via Sanger sequencing and Tracking of Indels by Decomposition (TIDE) analysis^33^. 20 candidate gRNAs were tested, and only five (E, I, F, J, 9) demonstrated the ability to edit the HIV-1 provirus within J-Lat cells (Figure 2B). Two of these gRNA target sites are located in the *gag* gene, within the p24 coding region (I, F); one spans the beginning of the *gag* gene including the first 10 bases of the p17 coding region; one gRNA target site is located in the Primer Binding Site sequence within the non-coding region of the virus (J); and one is located in the U5 region of the 5′ LTR (9) (Figure 2C). Additionally, these five gRNAs are all relatively conserved among HIV-1 M-group subtypes, and lack off-target sites within the human genome as determined by the off-target search tool, Cas-Offinder^34^ (Table 1). Overall, these five gRNA target sites represent ideal candidates for multiplex editing of HIV-1 M-group subtypes.

**Table 1.**
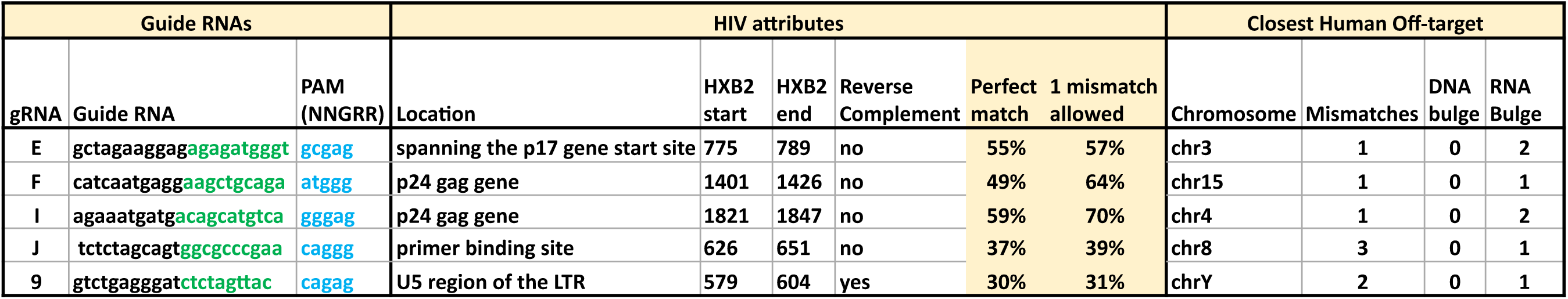
Characterization of HIV-1 gRNA target sites. gRNA sequence, location, HXB2 position start and finish, level of conservation among HIV-1 M-group subtypes (using an updated alignment from the 2020 HIV-1 database, that included 3,860 full length genomes), and the closest off-target sites are given. Exact match conservation corresponds to a perfect match with the entire 21 bp gRNA target site and the PAM site. One mismatch allows for a 1 bp mismatch to occur in the distal region (PAM-distal 11 nucleotides, bases in black lettering), but requires a perfect match in the 10 bases proximal to the PAM site (green lettering), and that the PAM motif be conserved. Chromosome location, nucleotide mismatch, and insertions (DNA bulge) or deletions (RNA bulge) are given for closest human genome off-target sites.

**Figure 2.**
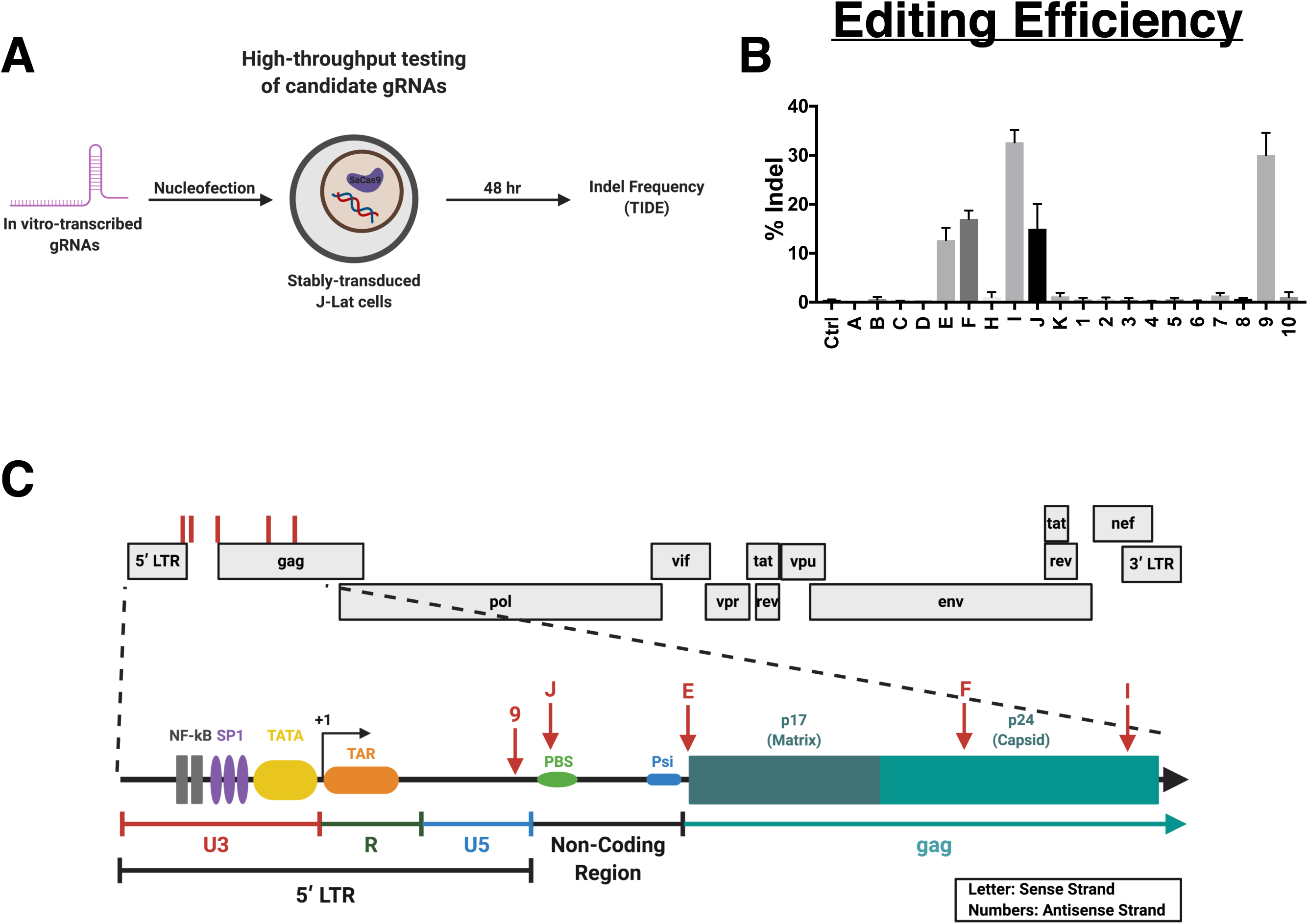
High-throughput screening of candidate HIV-1 gRNA target sites. (A) Workflow schematic of high-throughput testing of candidate HIV-1 gRNAs. *In vitro*-transcribed gRNAs were nucleofected into J-Lat cells stably-expressing SaCas9 and assessed for indels 48hr post-transfection. (B) Indel frequency of the five gRNAs capable of inducing SaCas9-mediated cleavage of latent provirus as measured by Sanger sequencing and TIDE analysis. (C) Schematic representation of the location of the five gRNAs capable of inducing indels within the latent provirus.

### Proviral editing results in diminished virion production and progeny infectivity

To accurately assess the ability of our gRNAs to edit HIV-1 provirus in latent cells, *in vitro*, and the subsequent effects on viral replication, we generated an HIV-1 latency model capable of producing infectious virions. We modified J-Lat cells to co-express HIV-1 HXB2 (CXCR4-tropic) *env* via lentiviral transduction followed by puromycin selection. The resulting J-Lats with Env provided *in Trans (*J-Lat-EnTr) cells constitutively express Env protein, and upon latency reversal are capable of producing single-round infectious virions with a GFP reporter in place of *nef* (Supplementary Figure 2A). J-Lat-EnTr cells expressed Env at levels comparable to Jurkat cells infected with WT HIV-1 (Supplementary Figure 2B). Additionally, GFP expression, before and after TNF-α stimulation, was comparable to that of the parental J-Lat cells (Supplementary Figure 2C). Finally, to assess the infectivity of virions produced from TNF-α-stimulated J-Lat-EnTr cells, viral supernatant was used to infect Jurkat cells in the presence or absence of the Integrase inhibitor, Raltegravir. Infectivity, assessed 3-days post-infection via GFP expression, showed that the parental J-Lat cells did not produce any infectious virus, whereas the J-Lat-EnTr cells produced virus capable of infecting Jurkat cells (Supplementary Figure 2D). Importantly, this infection could be blocked by Raltegravir, suggesting that GFP expression was produced via new infection as opposed to the possible contamination of GFP^+^ cells after TNF-α stimulation. These data suggest that the J-Lat-EnTr cell line can be used to accurately assess editing efficiencies of our gRNAs, and the subsequent effects on viral replication.

Next, we wanted to assess the ability of these five gRNAs to edit latent provirus when expressed from a lentiviral vector. Each individual gRNA was cloned into a novel lentiviral construct, pLenti-SaCas9-Neo, that is capable of simultaneously expressing SaCas9, a neomycin selection marker, and gRNA (Figure 3A). Virus-like particles (VLPs) containing each gRNA were produced and used to transduce J-Lat-EnTr cells. After neomycin selection, transduced cells were treated with TNF-α for 48hr, and subsequently assessed for editing efficiency via TIDE analysis, viral reactivation via J-Lat-EnTr GFP expression, virion production via p24 ELISA, and infectivity of progeny virus via target Jurkat GFP expression (Figure 3B). Editing efficiency was highest in cells transduced with gRNA I (72% Indels), followed by gRNA E (51% Indels) and gRNA F (43% Indels), indicating that targeting of the HIV-1 *gag* gene was most efficient (Figure 3C). Unfortunately, gRNA J (12% Indels) targeting the critical and extremely conserved PBS was associated with the lowest editing efficiency. Across tested gRNAs, editing efficiency was not affected by TNF-α stimulation, suggesting that reactivation of the latent provirus had no effect on editing efficiency (Figure 3C). As expected based on the absence of gRNAs targeting the viral LTR region, editing of the HIV-1 provirus by each gRNA had little to no effect on transcriptional reactivation as measured by GFP expression following TNF-α stimulation (Figure 3D). However, proviral editing of the *gag* gene (gRNAs I, E, F) exerted significant effects on virion production following TNF-α stimulation as measured by p24 ELISA of culture supernatants (Figure 3E). gRNA I (P < 0.0001) reduced virion production to undetectable levels at the dilution measured (1:100,000), while gRNA E (P < 0.0001) and gRNA F (P = 0.0038) reduced virion production by 4-fold and 2-fold, respectively, compared to cells transduced with SaCas9 alone. gRNAs J and 9 had no effect on virion production. Viral supernatant from TNF-α stimulated cells was used to infect Jurkat cells in culture, and infectivity was measured by GFP expression 72hrs post infection. gRNAs targeting the *gag* gene had the most significant effect on progeny virus infectivity, with gRNAs I, E, and F reducing infection by 8-fold (P =0.0002), 4-fold (P = 0.0005), and 2-fold (P = 0.0018), respectively (Figure 3F). gRNAs J (P = 0.0067) and 9 (P = 0.0047) were less effective but also reduced infectivity by statistically significant levels. Overall, these data show that CRISPR/Cas9 targeting of the HIV-1 *gag* gene is a highly effective means of reducing virion production and infectivity of new target cells, while the high sequence conservation of the targeted *gag* regions ensures efficacy across HIV-1 M-group strains.

**Figure 3.**
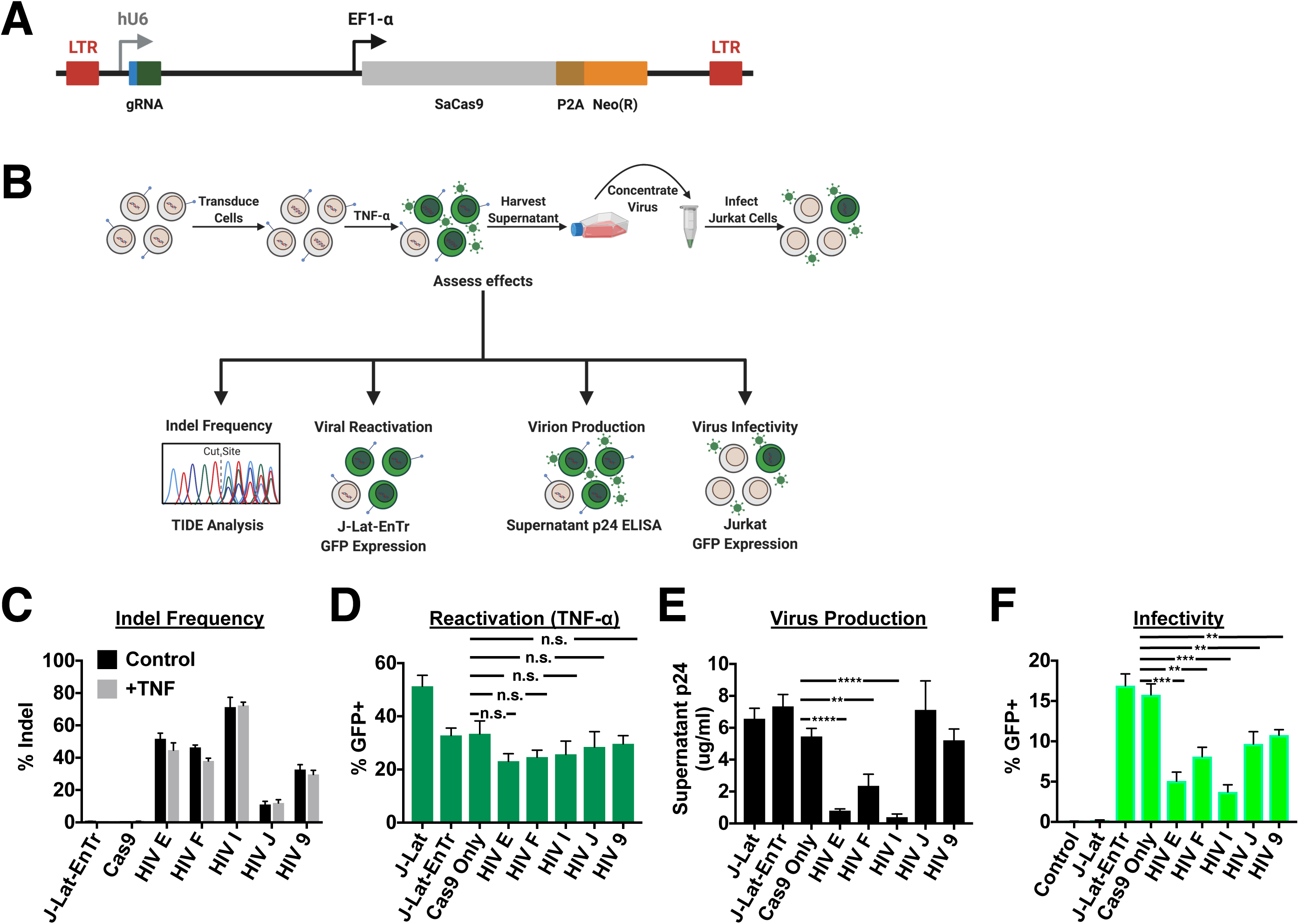
Editing efficiency of highly conserved HIV-1 gRNAs and their effects on virus replication in J-Lat-EnTr cell line. (A) Schematic of pLenti-SaCas9-Neo lentiviral vector used to express individual HV-1 gRNAs from the human U6 (hU6) promoter, and SaCas9 from an EF1-α promoter. A neomycin-resistant gene is also expressed from the EF1-α promoter via a P2A cleavage sequence. (B) Experimental workflow for assessing editing efficiency and the subsequent effects on viral replication. J-Lat-EnTr cells were transduced with virus-like particles (VLPs) containing individual gRNAs, selected via neomycin-resistance, and assessed for indels. Transduced cells were then stimulated with TNF-α for 48hr, and GFP expression was assessed via FACS. Supernatant was harvested, concentrated, and used to assess virion production via p24 ELISA. Supernatant was used to infect Jurkat cells, and GFP expression of Jurkat cells was assessed 72hr post infection. (C) Indel frequency of individual gRNAs +/- TNF-α treatment, and their effects on (D) transcriptional reactivation, (E) virion production, and (F) infectivity of progeny virus. The assays were performed in triplicate and error bars represent SD. Statistical significance was determined using unpaired t-tests, comparing each gRNA to Cas9 only control (\*\**P* < 0.01, \*\*\**P* < 0.001, \*\*\*\**P* < 0.0001, n.s. not significant).

### Co-targeting of latent provirus and CXCR4 reduces infection of new cells

To target the provirus at multiple sites, in order to prevent the emergence of escape mutants, we next combined our four best gRNAs (E, F, I, 9) for multiplex expression from our pLenti-SaCas9-Neo vector. This redundancy meant that 97% of the M group sequences matched at least one gRNA+PAM sequence, 74% matched at least 2, 39% matched 3, and 12% matched all 4, thus greatly enhancing the opportunity for editing diverse HIV-1 genomes at the population level and at the host quasispecies level relative to using any single gRNA (Supplementary Figure 3A).

First, we evaluated the use of different Pol III promoters for the expression of our gRNAs. Two separate lentiviral vectors were constructed: one with all four gRNAs being expressed from four separate human U6 promoters (hU6), and one with each gRNA being expressed from a distinct Pol III promoter (Supplementary Figure 4A). For this vector, we utilized the hU6 promoter, as well as the human H1 and 7SK promoters, and the murine U6 (mU6) promoter to drive expression of our four gRNAs targeting the HIV-1 provirus^35^. Transduction of J-Lat cells revealed that the use of distinct promoters produced a greater frequency of indels as opposed to the use of four separate hU6 promoters (Supplementary Figure 4B). Therefore, going forward, distinct Pol III promoters were used for multiplex expression of our gRNAs.

Next, we engineered a potential therapeutic CRISPR/Cas9 lentiviral vector that targets CXCR4 in uninfected target cells, rendering them non-permissive to HIV-1 infection, while simultaneously editing latent provirus in HIV-infected cells. In order to target both provirus and CXCR4 simultaneously, while guarding against viral escape mutants, we chose to express HIV-1 gRNAs I and F, along with a single gRNA targeting CXCR4 from pLenti-SaCas9-Neo (Figure 4A). gRNAs I and F had the greatest coverage potential, with 92% of HIV-1 sequences matching one of the two gRNAs, and 41% matching both (Supplementary Figure 3B). To test the potential of our therapeutic vector, we developed a co-culture assay consisting of virus-producing J-Lat-EnTr cells that express BFP (J-Lat-EnTr-BFP) in place of the *nef* ORF, and Jurkat target cells that constitutively express a GFP transgene from an EF1-α promoter (Jurkat-GFP). J-Lat-EnTr-BFP cells were generated from J-Lat-EnTr-GFP cells by creating a 194C > G and a 196T > C substitution using CRISPR-mediated homology-directed repair (HDR)^36^ (Supplementary Figure 5). A 2:1 ratio of Jurkat-GFP:J-Lat-EnTr-BFP co-culture was transduced with our therapeutic lentiviral vector and selected via neomycin-resistance (Figure 4B). Neomycin selection generated a pure population of transduced cells and allowed for the unbiased comparison of editing efficiencies and their subsequent effects among the different cellular populations. The transduced co-culture was then treated with a single dose of TNF-α for a period of 72hrs, during which time J-Lat-EnTr-BFP cells could produce single-round infectious virions capable of infecting Jurkat-GFP target cells with a readout of GFP/BFP double-positive cells being indicative of new infection events. Assessment of indel frequency for the entire co-culture, prior to TNF-α stimulation, revealed that our therapeutic vector could induce editing efficiencies of 29% and 32% at proviral target sites I and F, respectively, while CXCR4 was edited at 17% efficiency (Figure 4C). These editing efficiencies were lower than that of vectors targeting either provirus or CXCR4 alone. The lower editing efficiency of CXCR4 from our therapeutic vector resulted in a 35% loss of CXCR4 protein on the surface of Jurkat target cells compared to a 70% loss when targeting CXCR4 alone (Supplementary Figure 6A, B). Additionally, targeting provirus alone resulted in a significantly greater loss of virion production than targeting provirus with our therapeutic vector with virion production reduced by 85% and 78%, respectively (Supplementary Figure 6C, P = 0.0161). Despite the lower editing efficiencies of provirus and CXCR4, and the resulting effects on virion production and CXCR4 surface expression, the combination of targeting both using a single lentiviral vector resulted in a more substantial decrease in infected Jurkat target cells compared to targeting either alone (Figure 4D). While targeting either provirus or CXCR4 alone resulted in greater than 50% reduction in Jurkat target cell infection, targeting both provirus and CXCR4 resulted in an additional 15% reduction in infected cells, a statistically significant increase in efficacy (Figure 4E, P = 0.0004). Based on our mathematical model, the efficacies achieved by our dual-targeting vector are predicted to offer 513 days of drug-free remission in the absence of ART, and 591 days of remission following ART cessation, as compared to only a 9-day remission period following ART cessation in the absence of any intervention. These data recapitulate and reinforce the results of our viral dynamic model suggesting that targeting both HIV-1 provirus and co-receptor simultaneously in infected and susceptible cells results in greater protection against new infection events.

**Figure 4.**
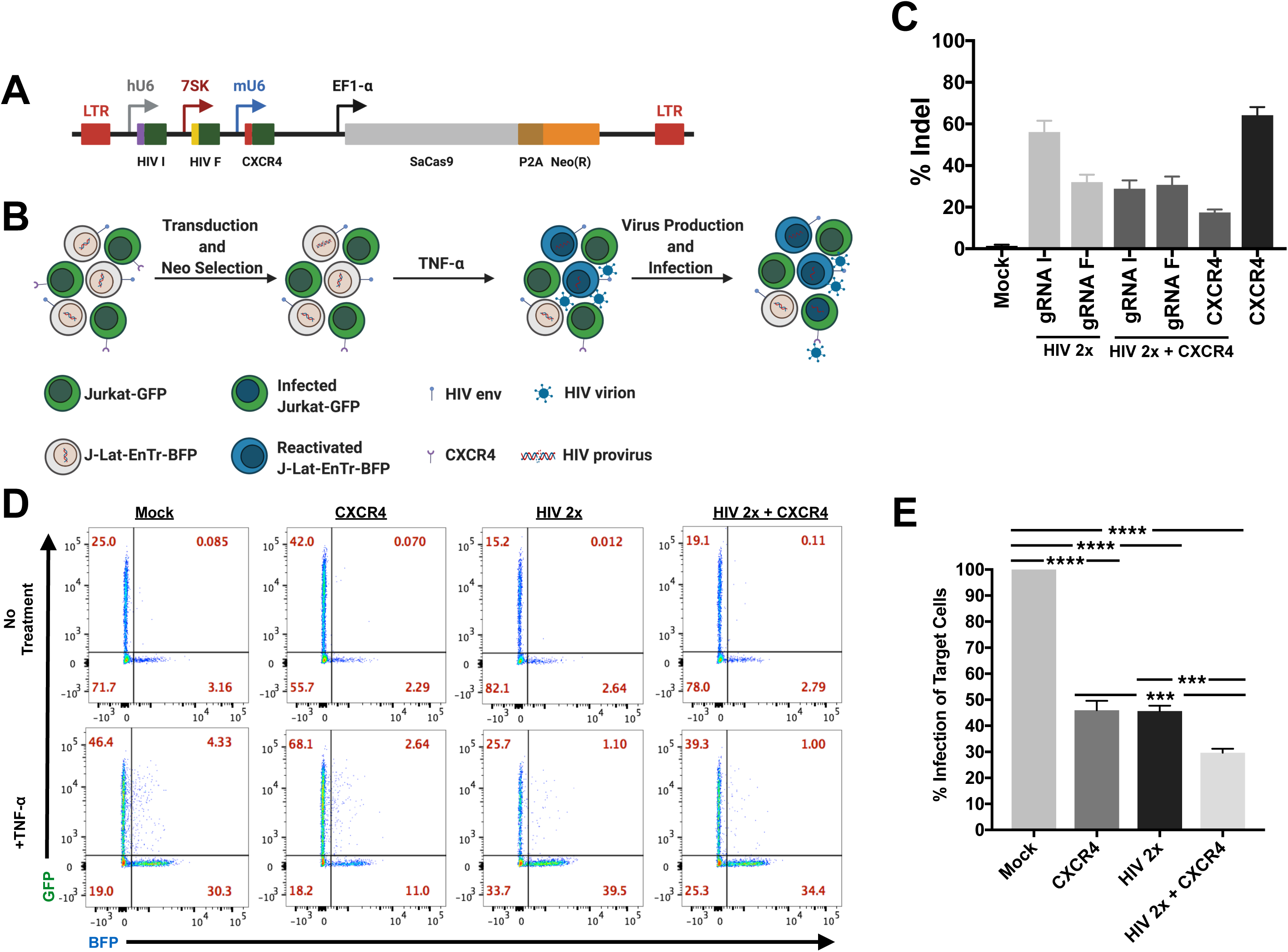
Targeting of latent provirus and CXCR4 in a co-culture assay. (A) Schematic representation of pLenti-SaCas9-Neo lentiviral vector used to express two gRNAs targeting HIV-1 and one gRNA targeting CXCR4. (B) Workflow of co-culture assay. Jurkat-GFP and J-Lat-EnTr-BFP cells were co-cultured at a 2:1 ratio and transduced with pLent-SaCas9-Neo multiplex vector. Transduced cells were then reactivated via TNF-α stimulation, and infectivity was assessed 72hr later via BFP^+^/GFP^+^ double-positive cells. (C) Indel frequencies at each target site. (D) Infectivity of Jurkat-GFP cells as assessed via GFP^+^/BFP^+^ double-positive cells pre and post TNF-α stimulation. FACS plots are representative of three independent experiments. (E) Quantified values for Jurkat-GFP cell infectivity from panel D. Experiments were performed in triplicate and error bars represent SD. Statistical significance was determined using One-way ANOVA and Bonferroni’s post-hoc tests for multiple comparisons, and unpaired t-test were used for comparisons between two groups (\*\*\**P* < 0.001, \*\*\*\**P* < 0.0001).

## DISCUSSION

The rapid advancement of CRISPR/Cas9 technology has led to the development of potential anti-HIV-1 gene editing therapeutic approaches. Thus far, these approaches have focused on targeting either the HIV-1 provirus or its co-receptors, CCR5 and CXCR4, with the majority of studies utilizing the *Strep. pyogenes* Cas9 (SpCas9) nuclease. Alone, these therapeutic approaches require editing efficiencies not yet achievable, *in vivo*^37,38^. However, deployed in a combinatorial approach, these two gene-editing strategies could potentially mimic the reduction in latent reservoir size (transplant conditioning) and the replacement of susceptible cells with resistant cells (CCR5^-/-^ transplant) that resulted in the only documented cases of HIV-1 cure. Therefore, our aim was to advance this therapeutic strategy by utilizing CRISPR/Cas9 technology to simultaneously edit latent provirus in infected cells and render uninfected cells refractory to infection. Using a novel *in vitro* co-culture assay, and an all-in-one lentiviral vector that expresses multiple gRNAs and the SaCas9 nuclease, we report the first proof-of-concept study supporting the two-pronged targeting approach of simultaneously editing provirus in infected cells and co-receptor in uninfected cells.

Previous clinical studies aimed at either reducing latent reservoir size or rendering cells refractory to infection have shown promise that a functional cure is possible, but have ultimately failed^39-42^. Reducing the reservoir size via a bone marrow transplant with uninfected CCR5^+^ (wildtype) donor cells afforded the “Boston patients” a brief remission period of 3-7 months before viral rebound. However, the pre-transplant conditioning failed to adequately deplete the latent reservoir, and the use of CCR5^+^ donor provided target cells capable of supporting viral rebound. Likewise, adoptive transfer studies of *ex vivo* CCR5-edited CD4^+^ T cells and hematopoietic stem cells proved safe in patients, but failed to achieve a remission period due to low editing efficiency and inefficient engraftment^37,38^. The failure of these single intervention approaches can be mitigated by a combinatorial approach.

Our mathematical modeling suggests that in order to achieve long-term remission, >88% of cells must be made co-receptor-deficient, or the latent reservoir must be reduced to <1% of its pre-intervention size. These numbers are in accordance with previous mathematical models^25^. However, previous studies have failed to adequately address the synergistic effect of combined interventions. Based on the successful combinatorial approach used with the Berlin and London patients, we have calculated that long-term remission is achievable, in the presence or absence of ART, by reducing the latent reservoir to below 40% of its original size and making 68% of uninfected cells HIV-resistant.

We engineered an all-in-one lentiviral vector capable of expressing two gRNAs targeting HIV-1, a single gRNA targeting CXCR4, and the SaCas9 enzyme. The use of SaCas9 instead of the more popular SpCas9 frees up ∼1kb of space in the lentiviral vector and increases the packaging capacity for gRNAs. The two HIV-1 gRNAs target highly conserved regions of the p24 capsid protein within the *gag* gene, and combined, are capable of targeting 92% of M-group subtypes. Additionally, these two gRNAs share negligible sequence homology with the human genome. This limits the potential for off-target editing associated with constitutive expression from lentiviral vectors, while allowing us to target multiple subtypes and guard against escape mutants. Additionally, the expression of all three gRNAs from distinct Pol III promoters as opposed to using multiple U6 promoters likely reduced the potential of recombination resulting from multiple repeated promoters that can cause the deletion of one or multiple cassettes^43^. This strategy resulted in improved editing efficiencies. This strategy could be further optimized and de-risked for clinical application by developing inducible Pol III promoters which restrict HIV-1 gRNAs expression to HIV-1-infected cells^44-47^.

We used a novel *in vitro* co-culture assay to show that our all-in-one lentiviral vector was capable of editing latent provirus in infected cells, and CXCR4 in uninfected cells. Despite exhibiting lower editing efficiencies than the single-targeting approaches, this dual-targeting approach reduced infection within our co-culture assay more significantly than either single-targeting approach. Proviral editing resulted in a 78% reduction in virion production, while co-receptor editing resulted in a 35% loss of surface expression on target cells. Our *in vitro* editing of provirus and co-receptor via our all-in-one lentiviral vector approached but did not reach the thresholds of our mathematical modeling that would result in indefinite remission. However, if achieved *in vivo*, these efficacies are predicted to offer 513 days of drug-free remission in the absence of ART, and 591 days of remission following ART cessation, as compared to only a 9-day remission period following ART cessation in the absence of any intervention. Further, subsequent rounds of gene therapy could be administered in order to reach the thresholds required to achieve indefinite remission. Alternatively, this dual-targeting approach could be used in combination with other cure strategies. Overall, these results underscore the potential of such a combinatorial approach in future therapeutic interventions.

In conclusion, this study presents the first proof-of-concept use of a bispecific gene editing approach to achieve a functional cure in HIV-1 infected individuals. Our *in silico* and *in vitro* data present a strong case for the use of this approach in future therapeutic interventions and warrant further investigation *in vivo*.

## METHODS

### Model of dual gene therapy: targeting host cell susceptibility and virion production

We model the effect of a dual-targeting gene therapy intervention which causes virion production in infected CD4^+^ cells to be reduced by a fraction, *f*_*v*_, and the susceptible pool of CD4^+^ cells to be reduced by a fraction, *f*_*s*_. We adopt a modified standard model of in host HIV-1 infection dynamics from (25) using parameters and state variables described in (25, 29-31).

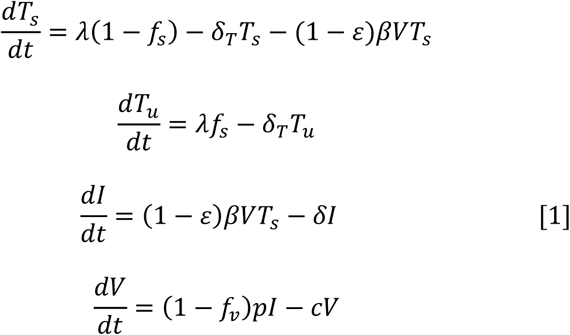

Where *T*_*s*_, *T*_*u*_, *I*, and *V* are susceptible target cells, refractory (non-susceptible) target cells, infected cells, and free virus, respectively. Parameters and initial conditions are consistent with the standard model of in host HIV-1 infection from (29) and (30). Uninfected target cells are produced at rate λ and have a death rate δ_T_. Free virus is produced at rate *p* and cleared at rate *c*. Susceptible cells are infected at rate β and have a death rate of δ. The fraction of viral production in infected CD4^+^ cells reduced by antiretroviral therapy (ART) is defined as ε. Assuming no intervention (*f*_*v*_ = *f*_*s*_ = 0) and ART (ε = 0), the reproductive number (*R*_0)_ for infected cells as described^30^ is:

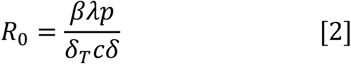

Considering the case where gene therapy modulates the fraction of CD4^+^ cells susceptible to infection and/or virion production in infected cells, we can define the reproductive number after gene therapy 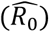 as:

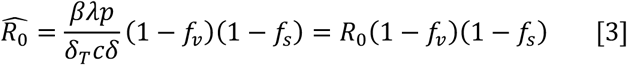

Using reproductive number *R*_0_ = 8 as previously estimated^31^, we can estimate the minimum efficiency of gene therapy needed to achieve a viral load at equilibrium of zero (durable control of viremia) as:

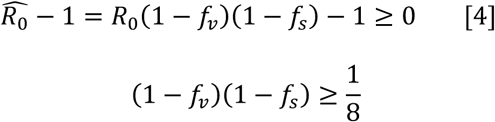

For gene therapy that reduces the fraction of susceptible CD4^+^ cells but does not affect virion production (*f*_*v*_ = 0), if >88% of target cells become refractory to infection, the model suggests that such intervention can lead to durable control, consistent with previous estimates^25^. However, with gene therapy that can both cause a reduction in the fraction of susceptible CD4^+^ cells and a reduction in viral production from infected cells, the threshold to achieve durable control is lowered. For example, interventions that reduce viral production by 60% and the fraction of susceptible cells by 68% can still lead to durable control.

To estimate the ART-free remission period after gene therapy, we use equation [1] to determine time *t* after cessation of ART (ε = 0) to reach clinically detectable viremia (*V*(*t*) > 50) for Figure 1. We assume ART is effective at suppressing all viral production during treatment (ε = 1) and that gene therapy is administered during continuous ART. *V*(*t*) is allowed to reach a steady state under ART prior to cessation of treatment. If *V*(*t*) < 50 for 100 years after cessation of ART, we consider the gene therapy to lead to durable control of viremia. A modified method was used to measure remission period using gene therapy in the complete absence of ART treatment (ε = 0). The remission period in the complete absence of ART was considered as the duration after administration of gene therapy with viremia below clinically detectable levels (*V*(*t*) < 50) for the analysis in Supplementary Figure 1.

### HIV-1 proviral gRNA target design

To design conserved gRNAs for the *Staph. aureus* Cas9 (SaCas9), we specified a gRNA length of 21bp, followed by the PAM sequence: NNGRR. These designs were generated in 2017, using the HIV-1 Los Alamos database filtered full-length genome set which included one sequence per sampled individual, filtered for high quality complete sequences with minimal ambiguity codes. The alignment included 3,263 sequences and was based on the global M group; we used the 2020 alignment that included 3,860 sequence to generate the coverage data in Table 1 and Supplementary Fig. 3A, B. We made a reverse complement set as well, to increase the opportunity to identify conserved gRNAs. We required that gRNAs did not have strings of four or more T’s (TTTT), as this might result in premature transcriptional termination of gRNAs from a Pol III promoter. We favored guide RNAs with 40-60% GC content, and avoided long strings of the same nucleotide.

### SaCas9-gRNA plasmid construction

pLenti-SaCas-Puro was constructed from the pSicoR-mCherry backbone. The U6 promoter was replaced with a U6 promter-SaCas9 gRNA scaffold gBlock (Integrated DNA Technologies (IDT)) containing two BsmBI restriction sites downstream of the U6 promoter using the XbaI and XhoI restriction sites. mCherry was replaced with SaCas9-P2A-Puro using NheI and EcoRI. A unique BstBI restriction site was included between SaCas9 and the P2A sequence. Lastly, bovine growth hormone and SV40 polyadenylation signals were cloned downstream of the 3’ LTR. pLenti-SaCas9-Neo was created by replacing with the Puro(R) gene with a Neo(R) gene using the BstBI and EcoRI restriction sites. Individual gRNAs were cloned into pLenti-SaCas9-Neo utilizing the two BsmBI restriction sites. For multiplex gRNA expression, promoters and gRNAs were synthesized as gBlocks (IDT) and cloned into pLenti-SaCas9-Neo via Golden Gate assembly utilizing the two BsmBI restriction sites.

### High-throughput testing of candidate gRNAs targeting HIV-1 provirus

J-Lat clone 11.1 cells (a kind gift from Eric Verdin) were transduced with pLenti-SaCas9-Puro and selected with 1ug/ml puromycin (Lifetech) to generate SaCas9 stably expressing J-Lats. *In vitro* transcription was used to synthesize candidate gRNAs. Primers were used to PCR amplify the SaCas9 gRNA scaffold from pLenti-SaCas9-Puro. All forward primers included a T7 promoter and individual gRNA target sequence upstream of the gRNA scaffold. PCR amplicons were used as template for *in vitro* transcription via T7 polymerase (New England BioLabs (NEB)). Transcribed gRNAs were treated with DNAse and purified using RNA Clean and Concentrator-100 (Zymo Research). gRNAs were transfected into SaCas9 expressing J-Lat 11.1s using the Amaxa 4D-Nucleofector X Unit (Lonza) and the Amaxa 4D-Nucleofector Protocol for Jurkat clone E6.1. PCR amplicons corresponding to each gRNA target region were amplified from genomic DNA isolated from cells 48hr post transfection, and subjected to Sanger sequencing and subsequent TIDE analysis.

### Generation of J-Lat-EnTr and J-Lat-EnTr-BFP cell lines

To generate J-Lat-EnTr cells, we PCR amplified HXB2 *env* from the pIIIenv3-1 plasmid (NIH AIDS Reagent Program cat. #289) and cloned it into the pLVX-EF1α-IRES-Puro lentiviral vector (Clontech cat. #631988). J-Lat clone 11.1 cells were transduced with lentiviral particles produced from this plasmid and stable cell lines were generated using puromycin selection.

To generate J-Lat-EnTr-BFP cells, we adapted a previously described method to convert GFP to BFP using CRISPR-Cas9 technology^36^. The eGFP sequence from the parental J-Lat-EnTr cell line was modified using a CRISPR-Cas9 RNP (ribonucleoprotein) mediated HDR approach. Synthetic crRNA targeting *EGFP* nucleotides responsible for eGFP fluorescence (5’CTCGTGACCACCCTGACCTA) and tracrRNA (IDT) were incubated at 37°C for 30 minutes to form an 80 μM guide RNA. Guides were then incubated with an equal volume of 40μM Cas9-NLS purified protein (UC QB3-Berkeley Macrolab) at 37°C for 15 minutes to form RNPs at 20μM. RNPs were then nucleofected into 10^6^ J-Lat 11.1 using a Lonza 4D-Nucleofector along with 100pmole of single stranded DNA HDR control template (consisting of a 132bp scrambled sequence) or a BFP HDR template (Sequence: 5’CTGAAGTTCATCTGCACCACCGGCAAGCTGCCCGTGCCCTGGC CCACCCTCGTGACCACCCTGAGCCACGGGGTGCAGTGCTTCAGCCGCTACCCCGACCAC ATGAAGCAGCACGACTTCTTCAAGTCCGCC 3’) (IDT). Nucleofected cells were then put in culture for 48 hours, followed by a 24 hours TNF-α treatment (20ng/mL) to reactivate the provirus and get fluorescent reporter expression. BFP^+^GFP^-^cells were then sorted by FACS using a Sony M900 cell sorter. Sorted cells were maintained in culture for one week to let them go back to latency. Latent cells (BFP^-^/GFP^-^) were then sorted and an aliquot was reactivated as previously described to evaluate the purity of the sorting and the ability of the provirus to reactivate following TNF-α treatment.

### Cell treatment

Non-transduced J-Lat and J-Lat-EnTr cells, and transduced J-Lat-EnTr cells were treated with 20ng/ml TNF-α for 48hr. Co-cultures of transduced J-Lat-EnTr-BFP and Jurkat-GFP (GenTarget Inc) were treated with 20ng/ml TNF-α for 72hr. Jurkat cells infected with supernatant from TNF-α stimulated J-Lat and J-Lat-EnTr cells were treated with 30ug/ml Raltegravir (NIH AIDS Reagent Program) immediately following spinoculation. Puromycin selection and neomycin selection was achieved at 1ug/ml and 5ug/ml, respectively.

### Virus production

All virus-like particles were generated in HEK293T cells via calcium phosphate transfection using packaging plasmids psPAX2 (Addgene #12260) and pMD2.G (Addgene #12259). Replication-competent HIV-1 NLENG-IRES-GFP (a kind gift from Warner Greene) was generated in HEK293T cells using calcium phosphate transfection. VLP and HIV-1 supernatant was harvested 48hr post transfection and concentrated via ultracentrifugation. Single-round replication-incompetent HIV-1 produced from J-Lat-EnTr cells was harvested 48hr post TNF-α treatment and concentrated via Lenti-X-Concentrator (Clontech). J-Lat-EnTr-BFP cells co-cultured with Jurkat-GFP cells produced single-round replication-incompetent HIV-1 produced via TNF-α treatment for 72hr. All virus supernatant was quantified via p24 ELISA (Perkin Elmer).

### Cell infection and transduction

All VLPs were used to transduce the various cell lines via spinoculation at a concentration of 1ug of p24 per 1×10^6^ cells. Virus and cells were centrifuged at 2350 rpm and 37**°**C for 2hr in a volume of ≤100 ul before being returned to culture. pLenti-SaCas9-Puro was used to generate J-Lat cells stably expressing SaCas9, and pLenti-SaCas9-Neo was used for all other CRISPR/Cas9 experiments. HIV-1 NLENG-1-IRES-GFP was used to infect Jurkat cells via spinoculation at a concentration of 100 ng of p24 per 1×10^6^ cells. Virus produced from non-transduced and transduced J-Lat-EnTr cells treated with TNF-α were used to infect Jurkat cells via spinoculation at a volume of 100ul per 5×10^5^ cells. Virus produced from transduced J-Lat-EnTr-BFP treated with TNF-α were allowed to infect Jurkat-GFP cells for 72hr without concentrating the virus.

### Flow cytometry

Non-transduced J-Lat and J-Lat-EnTr cells were assessed for GFP expression 24hr post TNF-α treatment. Jurkat cells infected with HIV-1 NLENG-1-IRES-GFP were assessed for GFP expression 48hr post infection. Transduced J-Lat-EnTr cells were assessed for GFP expression 24hr post TNF-α treatment. Jurkat cells infected with supernatant from TNF-α treated J-Lat-EnTr cells and Jurkats infected with HIV-1 were assessed for GFP expression 72hr post infection. J-Lat-EnTr-BFP and Jurkat-GFP co-cultures were assessed for GFP, BFP, and CXCR4 surface expression 72hr post TNF-α treatment by staining in FACS buffer (phosphate buffered saline supplemented with 2 mM EDTA and 2% FBS) with α-CXCR4-APC (eBiosciences). All cells were fixed in 1% paraformaldehyde prior flow cytometry. All data were collected on a FACS LSRII (BD Biosciences), and analyses were performed with FlowJo software (TreeStar).

### Western Blot Protein Analysis

Uninfected and HIV-1-infected Jurkat cells, along with untreated and TNF-α treated J-Lat and J-Lat-EnTr cells were lysed in radioimmunoprecipitation assay buffer (150 mm NaCl, 1% Nonidet P-40 (vol/vol), 0.5% AB-deoxycholate (vol/vol), 0.1% sodium dodecyl sulfate (SDS) (vol/vol), 50 mm Tris-HCl (pH 8), 1 mm DTT, and EDTA-free Protease Inhibitor (Calbiochem). Cell lysates were used for SDS-polyacrylamide gel electrophoresis (SDS-PAGE) immunoblotting analysis. The primary antibodies used were mouse monoclonal α-HIV-1 IIIB gp160 (NIH AIDS Reagent Program #1209) at 1:100; mouse monoclonal α-HIV p24 AG3.0 (NIH AIDS Reagent Program #4121) at 1:100; and mouse monoclonal α-GAPDH (Abcam ab8245) at 1:1000.

### Statistical Analysis

The data were analyzed using GraphPad Prism 7.0 software (La Jolla, CA) and presented as the standard deviation (SD) of three independent experiments. One-way ANOVA and Bonferroni’s post-hoc tests where used for multiple comparisons and unpaired t-test were used for comparisons between two groups as indicated. Significant differences were determined at *P* < 0.05.

## AUTHOR CONTRIBUTIONS

L.R.C designed the study, performed experiments, analyzed data and wrote the paper; N.R.R. designed and implemented the viral dynamic model and co-wrote the paper; M.S.B. developed the J-Lat-EnTr-BFP cell line; K.A.R., S.D., Z.Y.D, and H.S.S. performed experiments; J.T. and B.K. conducted bioinformatic analyses to design gRNAs and co-wrote the paper; S.K.P. designed the study, analyzed data and wrote the paper.

## ACKNOWLEDGMENTS

This study was supported by the National Institutes of Health grants R01 AI150449 and R01 MH112457 (to SKP), K12 HL143961 (to LRC), and was additionally supported by a grant from the National Institutes of Health, University of California San Francisco-Gladstone Institute of Virology & Immunology Center for AIDS Research (P30 AI027763).

## FIGURE LEGENDS

**Supplementary Figure 1.**
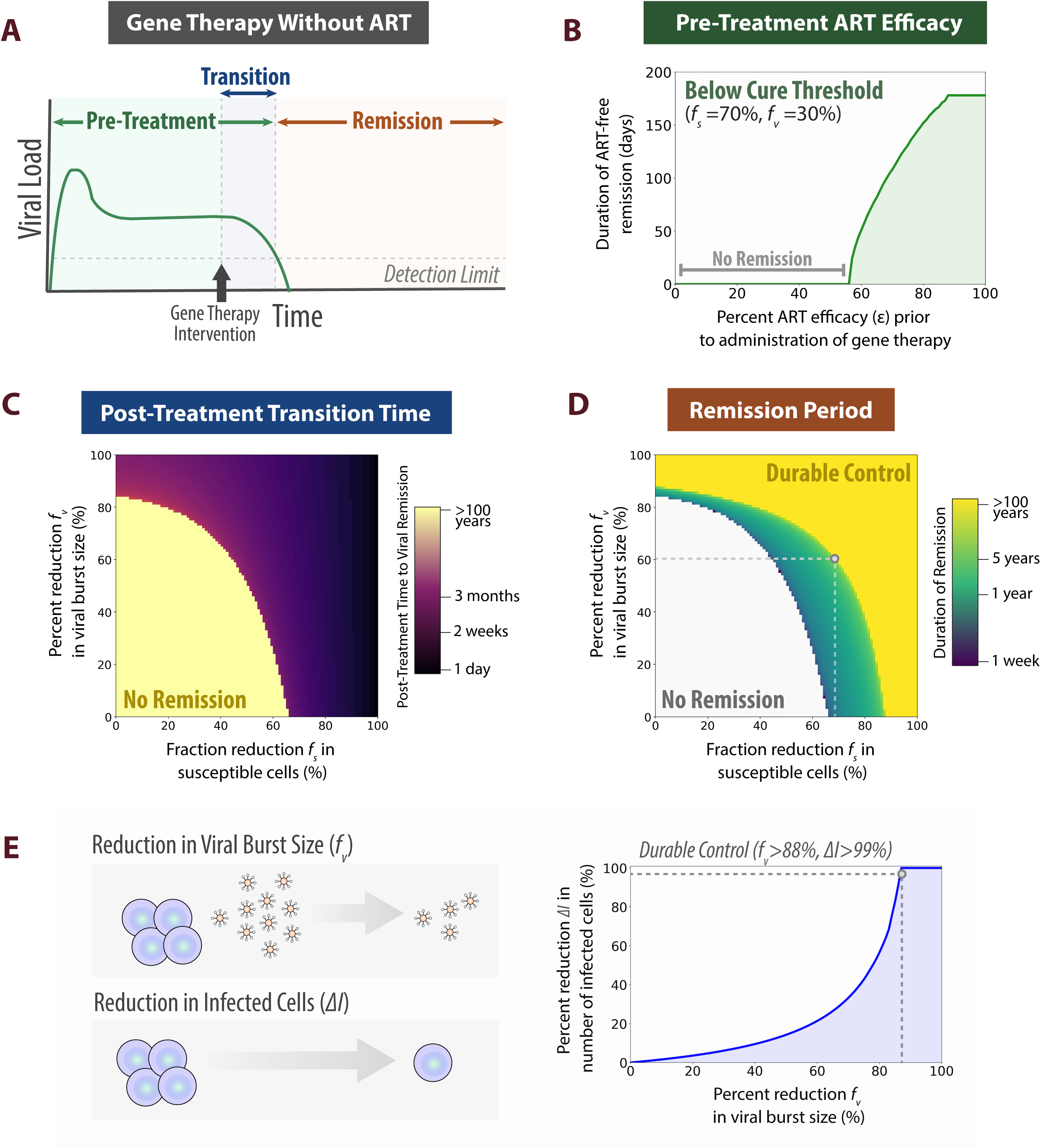
Computational predictions of within-host HIV-1 dynamics for bispecific gene therapy in the absence of antiretroviral therapy. (A) Gene therapy framework in the absence of antiretroviral therapy (ART). Gene therapy is administered after infection in the absence of ART, leading to remission if gene therapy is sufficiently efficacious. Post-treatment transition time describes the duration between administration of the gene therapy and reduction in the viral load to below the detection limit. A modified standard model of within-host HIV-1 dynamics is used to simulate this treatment strategy (see Methods). (B) The effects of pre-treatment ART efficacy on remission period of gene therapy below cure threshold efficacy. If the efficacy of gene therapy is below the cure threshold (ex. 70% reduction in cell susceptibility and a 30% decrease in virion production), the duration of the remission period is extended in proportion to ART efficacy. (C) Post-treatment transition time to suppress viral load to below detection limit. The efficacy of the bispecific gene therapy determines the time required to achieve viral remission. (D) Remission period with gene therapy in the absence of ART. The duration of remission post-treatment with gene therapy of varying efficacies are shown. When gene editing efficacy is low, viral load is not expected to drop below the detection limit and there is no remission period. High efficacy gene editing is predicted to lead to durable control (>100 years remission). (E) Relationship between viral burst size and number of infected cells. The percent change in the number of infected cells is shown in relation to the reduction in viral burst size after gene therapy intervention (cell susceptibility is constant, *f*_*s*_ = 0). A reduction in viral burst size by >88%, the threshold to achieve durable control, corresponds to a >99.9% reduction in the number of infected cells post-treatment.

**Supplementary Figure 2.**
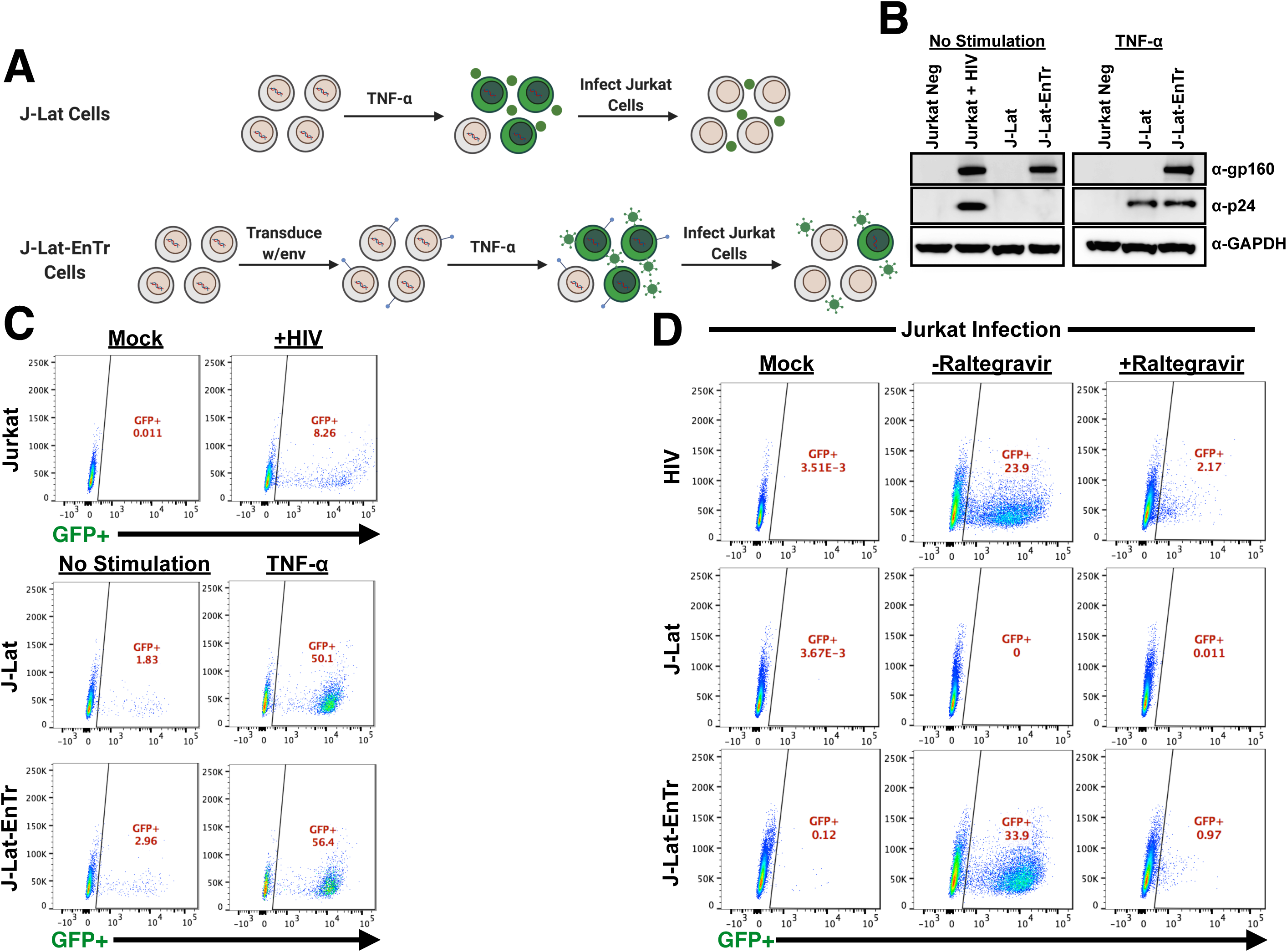
Development and characterization of J-Lat-EnTr cell line. (A) Generation of J-Lat-EnTr cell line. Parental J-Lat cells were transduced with VLPs containing cDNA for HIV-1 HXB2 Env under the control of EF1-α, and selected via puromycin resistance. When treated with TNF-α, J-Lat-EnTr cells express GFP, and generate virions containing env protein, while parental J-Lat cells only express GFP. Supernatant from J-Lat-EnTr cells can then be used to infect Jurkat cells, with a readout of GFP expression for infection. (B) Protein expression of HIV-1 gp160 and p24 in unstimulated and TNF-α stimulated J-Lat and J-Lat-EnTr cells. (C) GFP expression in unstimulated and TNF-α stimulated J-Lat and J-Lat-EnTr cells. (D) Infection of Jurkat cells using supernatant from TNF-α stimulated J-Lat and J-Lat-EnTr cells +/- the integrase inhibitor, Raltegravir. As a positive control, Jurkat cells were infected with WT HIV-1.

**Supplementary Figure 3.**
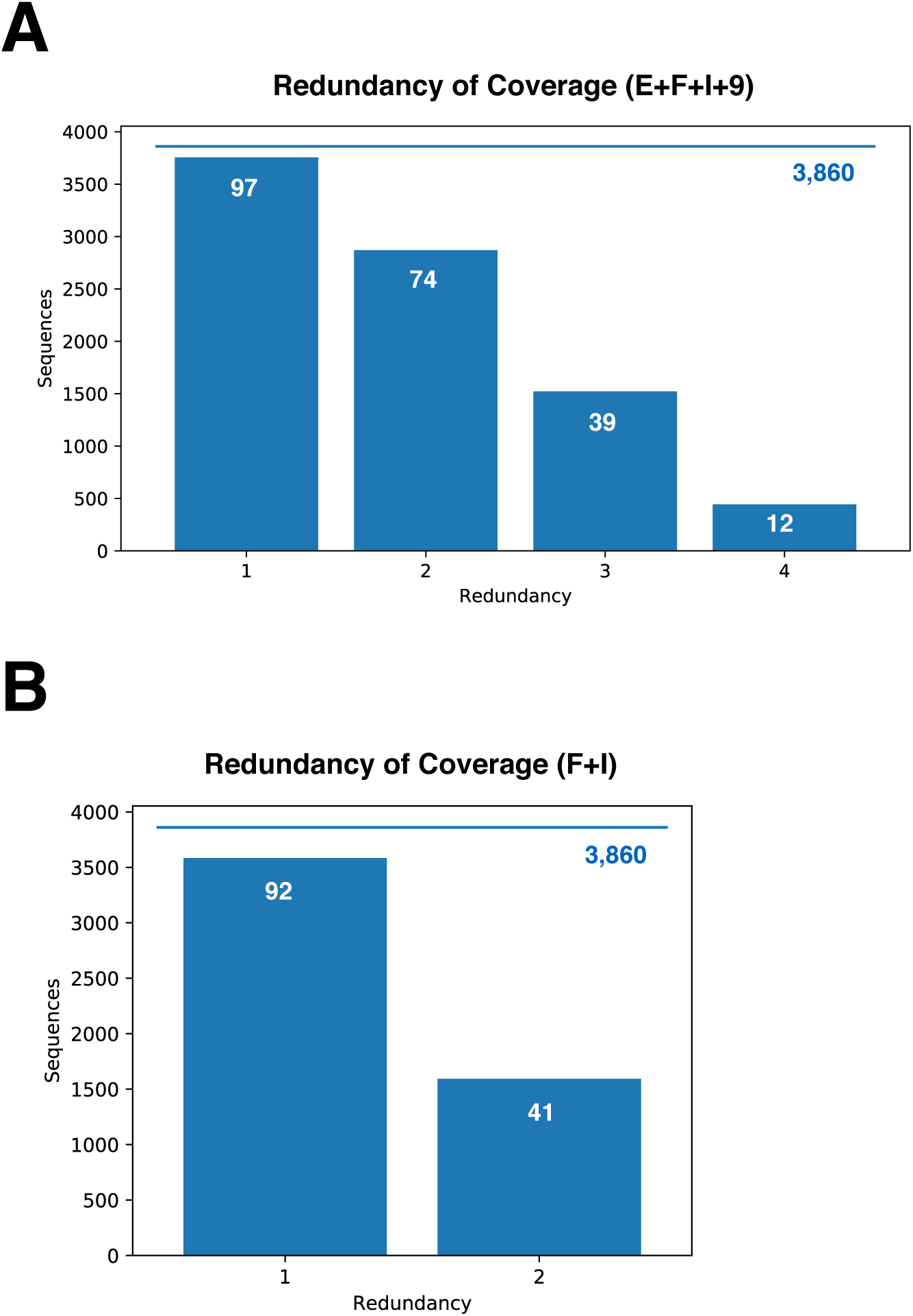
HIV-1 M group sequence redundancy of coverage for multiplexed gRNAs expressed from pLenti-SaCas9-Neo vector. (A) The enhanced sequence coverage for gRNA redundancy in the delivery of 4 gRNAs (E, F, I, and 9) from a single lentiviral vector. (B) The enhanced sequence coverage for gRNA redundancy in the delivery of the two best gRNAs (I and F) that were paired with the CXCR4 gRNA into a single lentiviral vector. Redundancy of coverage is calculated based on the 1-mismatch-allowed criteria. The percent M group coverage is written in white within each blue bar.

**Supplementary Figure 4.**
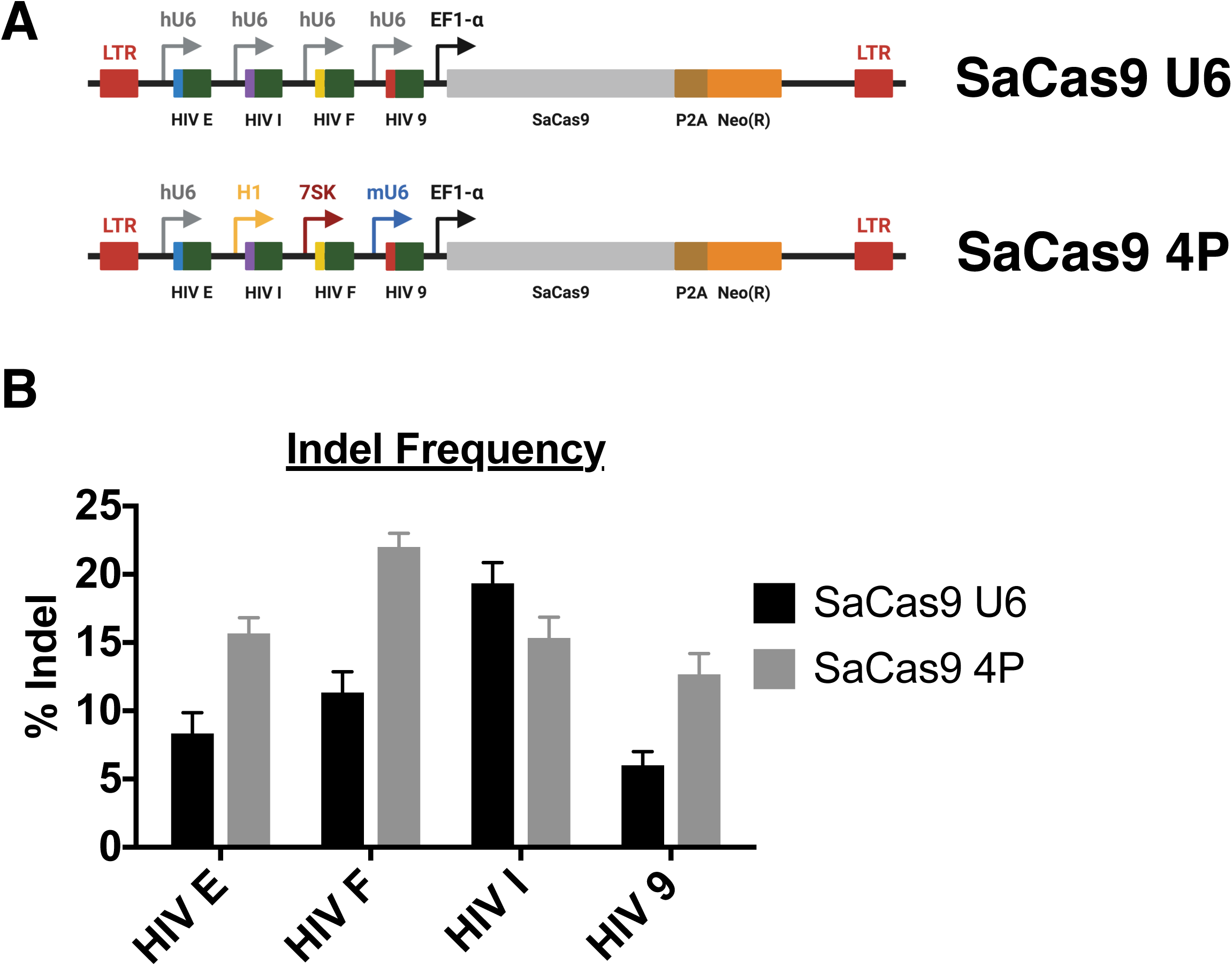
Comparison of Pol III promoters for the expression of gRNAs in multiplex lentiviral vector. (A) Schematic of pLenti-SaCas9-Neo vectors used to express multiple gRNAs using either four hU6 promoters or four distinct Pol III promoters. (B) Indel frequency of individual gRNAs in J-Lat-EnTr cells when expressed from multiplex vectors.

**Supplementary Figure 5.**
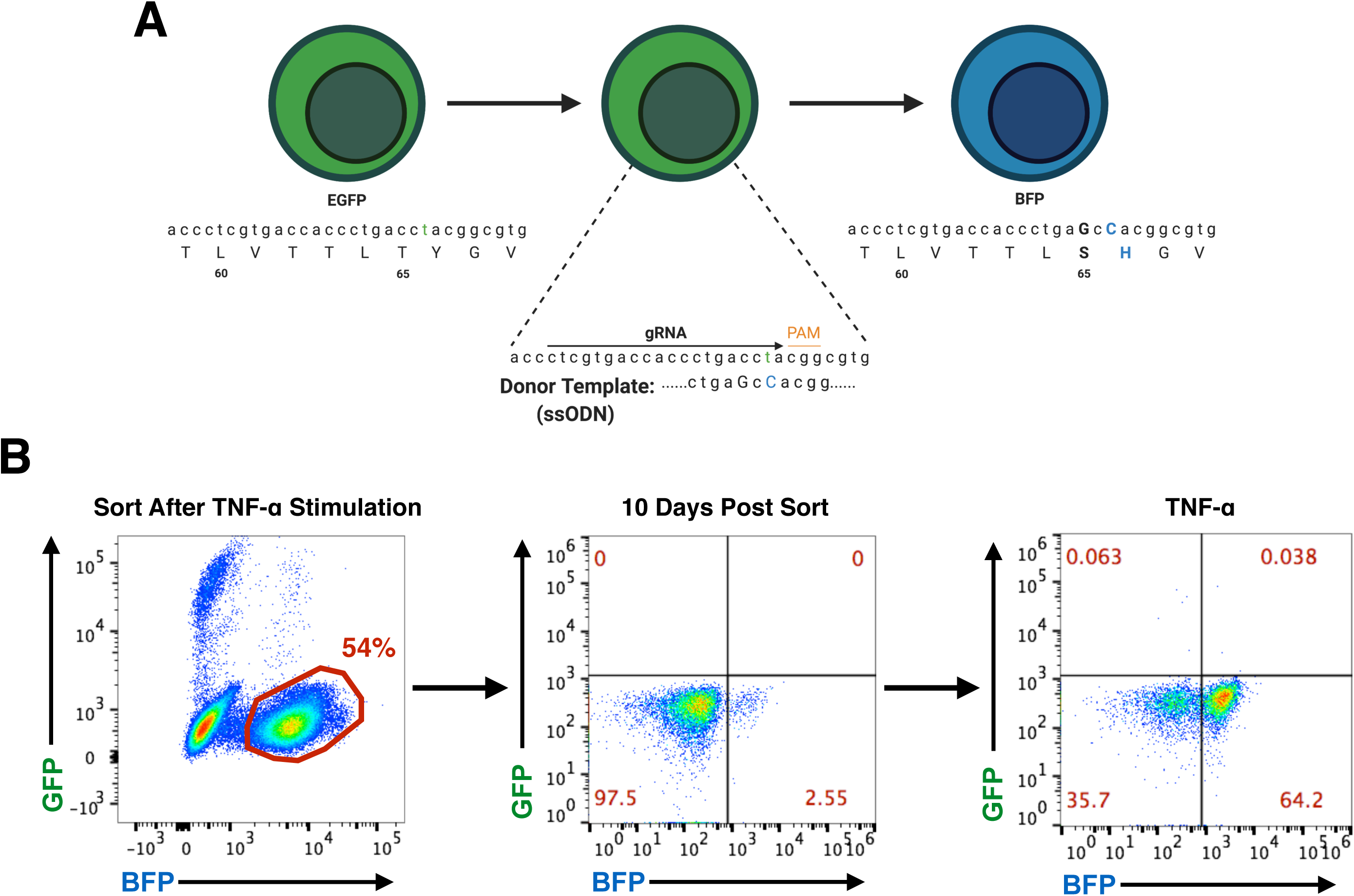
Generation of J-Lat-EnTr-BFP cell line using CRISPR/Cas9-mediated HDR. (A) Gene targeting strategy. A donor template containing a 194C > G and a 196T > C substitution was used to convert eGFP to BFP in J-Lat-EnTr cells. (B) Sort strategy and subsequent culture of J-Lat-EnTr-BFP cells. J-Lat-EnTr cells were transfected with gRNA-Cas9 RNP and donor template. BFP-positive cells were sorted 24hr post TNF-α treatment. Sorted cells were then returned to culture and gradually lost BFP expression over the course of 10 days. Subsequent stimulation with TNF-α revealed a pure J-Lat-EnTr-BFP population.

**Supplementary Figure 6.**
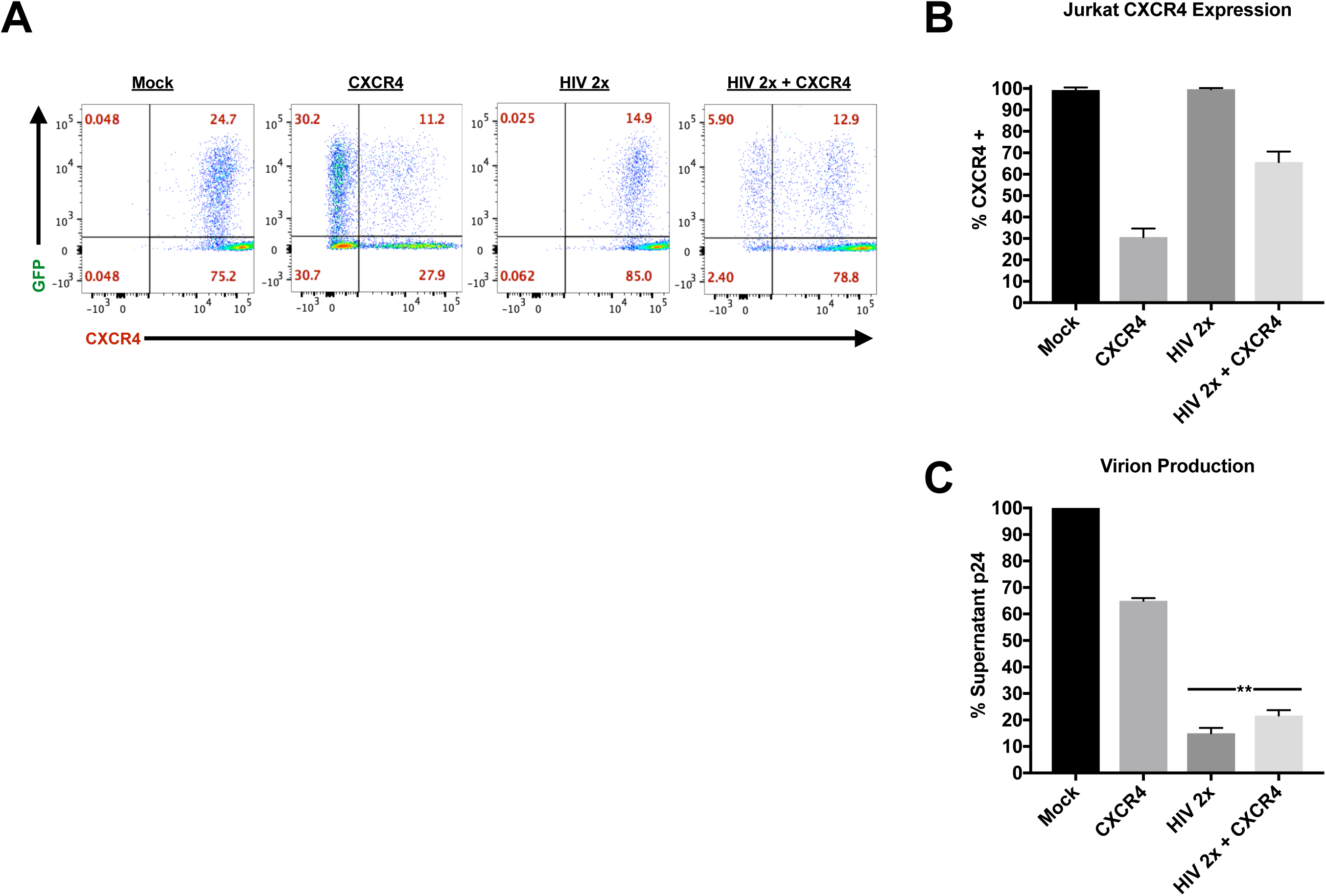
Effects of multiplex editing on CXCR4 surface expression and virion production. (A) CXCR4 surface expression on Jurkat-GFP target cells after SaCas9-mediated CXCR4 editing. FACS plots are representative of three independent experiments. (B) Quantified values for Jurkat-GFP surface expression of CXCR4 from panel A. (C) Virion production from transduced J-Lat-EnTr-BFP cells in co-culture with Jurkat-GFP cells 72hr post TNF-α stimulation. The assays were done in triplicate and error bars represent SD. Statistical significance was determined using unpaired t-test (\*\**P* < 0.01).

## Notes

**Conflict of interest statement** The authors have declared that no conflict of interest exists.

### Competing Interest Statement

The authors have declared no competing interest.

